# Upstream translation initiation expands the coding capacity of segmented negative-strand RNA viruses

**DOI:** 10.1101/795815

**Authors:** Elizabeth Sloan, Marta Alenquer, Liliane Chung, Sara Clohisey, Adam M. Dinan, Robert Gifford, Quan Gu, Nerea Irigoyen, Joshua D. Jones, Ingeborg van Knippenberg, Veronica Rezelj, Bo Wang, Helen M. Wise, Maria Joao Amorim, J Kenneth Baillie, Ian Brierley, Paul Digard, Andrew E. Firth, Megan K. MacLeod, Edward Hutchinson

## Abstract

Segmented negative-strand RNA viruses (sNSVs) include the influenza viruses, the bunyaviruses, and other major pathogens of humans, other animals and plants. The genomes of these viruses are extremely short. In response to this severe genetic constraint, sNSVs use a variety of strategies to maximise their coding potential. Because the eukaryotic hosts parasitized by sNSVs can regulate gene expression through low levels of translation initiation upstream of their canonical open reading frames (ORFs), we asked whether sNSVs could use upstream translation initiation to expand their own genetic repertoires. Consistent with this hypothesis, we showed that influenza A viruses (IAVs) and bunyaviruses were capable of upstream translation initiation. Using a combination of reporter assays and viral infections, we found that upstream translation in IAVs can initiate in two unusual ways: through non-AUG initiation in virally encoded ‘untranslated’ regions, and through the appropriation of an AUG-containing leader sequence from host mRNAs through viral cap-snatching, a process we termed ‘start-snatching.’ Finally, while upstream translation of cellular genes is mainly regulatory, for sNSVs it also has the potential to create novel viral gene products. If in frame with a viral ORF, this creates N-extensions of canonical viral proteins. If not, it allows the expression of cryptic overlapping ORFs, which we found were highly conserved in IAV and widely distributed in peribunyaviruses. Thus, by exploiting their host’s capacity for upstream translation initiation, sNSVs can expand still further the coding potential of their extremely compact RNA genomes.

## Introduction

Viruses operate under extreme coding constraints. Their small size, replication strategies and high replication error rates severely limit the total length of their genomes, particularly for the error-prone RNA viruses [1, 2]. As a result, viruses have evolved diverse strategies to increase their coding potential [1, 3]. For example the influenza viruses, whose RNA genomes are less than 14 kb in length, are known to encode alternative gene products through mechanisms including alternative splicing, ambisense coding, leaky ribosomal scanning, ribosomal frameshifting and termination-dependent re-initiation [4–7]. As viruses lack their own protein synthesis machinery, they must be able to encode all of their proteins in a form that host ribosomes will recognise as mRNA [8]. In eukaryotes, ribosomes recognise mRNAs with a terminal 5′ cap structure followed by an untranslated region (UTR), which can be tens to hundreds of nucleotides in length [9–11]. A growing body of work has shown that translation initiates at low levels in the 5′ UTRs of a large proportion of eukaryotic mRNAs, sometimes extremely close to the 5′ cap, and that translation of the resulting upstream open reading frames (uORFs) can regulate cellular gene expression [10,12–18]. As a result, the need to mimic their host’s mRNA structure could provide viruses with the potential to exploit upstream translation initiation, something which could allow them to expand the coding potential of their own genomes.

To determine whether this was the case, we examined segmented, negative-strand RNA viruses (sNSVs). These include the orthomyxovirus family (which includes the influenza viruses) and the order of bunyaviruses (which include the aetiological agents of haemorrhagic fevers and other severe human illnesses, as well as many other pathogens of humans, animals and plants). The sNSVs are a useful order of viruses in which to search for upstream translation initiation for three reasons. Firstly, as RNA viruses they are already known to employ a variety of alternative coding strategies in response to genetic constraints, which suggests that they may also have undergone selection for additional alternative coding methods. Secondly, each segment of an sNSV genome produces its own mRNAs with distinct 5′ UTRs, and this diversity of viral 5′ UTRs increased the likelihood that we might observe upstream translation initiation. Thirdly, sNSVs produce their mRNA through cap-snatching, a process in which 7-methylguanylate (m^7^G)-capped 5′ termini are cleaved from host mRNAs to provide RNA primers for the transcription of viral genes [11]. The length of cap-snatched host sequences varies, but cap-snatching typically results in 10 – 13 nt of host encoded-sequence being appended to the 5′ end of the mRNAs of IAV [19–22] and 11 – 18 nt to the mRNAs of the bunyaviruses [23–27]. In other words, as a result of cap-snatching sNSVs produce chimeric mRNAs in which host 5′ UTR sequences are present upstream of the virally-templated 5′ UTR. It seemed plausible that, if upstream translation initiation was occurring in host mRNAs, it might also occur in these chimeric viral mRNAs.

To determine whether upstream translation initiation could occur in sNSVs, we examined three viruses: the orthomyxovirus influenza A virus (IAV), a cause of influenza in humans and other animals and the cause of all influenza pandemics [28], and two bunyaviruses: the tick-borne phenuivirus Heartland virus [29] and the midge-borne peribunyavirus Oropouche virus [30]. We demonstrated that, consistent with our hypothesis, each of these viruses is capable of upstream translation initiation. Focussing on IAV, we showed that upstream translation initiation takes place through two distinct routes – initiation within the virally-encoded 5′ UTR, and initiation still further upstream in the host-derived cap-snatched leader. This second route produces chimeric gene products in a process we termed ‘start-snatching.’ We found that either route for upstream translation can, depending on viral sequence, produce N-terminal fusions to viral proteins, and we identify an example of such an N-terminally extended proteins within influenza virions. In addition, it is possible for upstream translation to access cryptic uORFs that overlap the canonical viral open reading frame (ORF). We discovered that the genetic potential to generate such uORFs is widely distributed in influenza viruses and in peribunyaviruses, and in the case of IAV we found evidence that uORFs are conserved, can be translated and can be presented to the adaptive immune system. We conclude that upstream translation initiation is a previously unappreciated mechanism for expanding the genetic capacity of sNSVs, increasing still further the ability of these viruses to encode a diverse array of gene products in a constrained genome.

## Results

### Upstream translation initiation occurs in segmented negative-strand viruses

We tested the hypothesis that upstream translation initiation could occur in sNSVs using minireplicon assays, transfection-based systems that generate reporter proteins through viral gene expression. In these assays the viral polymerase and nucleoprotein transcribe a negative-strand template RNA, consisting of a reporter gene flanked by viral UTRs, into an mRNA which is translated to give the reporter protein. We used minireplicon systems encoding luciferase (Luc) reporters to test the transcriptional machineries of three sNSVs – an orthomyxovirus (the influenza A virus (IAV) A/WSN/33(H1N1) (WSN); NS segment UTRs), a phenuivirus (Heartland banyangvirus (HRTV); L segment UTRs) and a peribunyavirus (Oropouche orthobunyavirus (OROV); M segment UTRs).

We first determined how much of the reporter signal was due to translation initiation at sites up to and including the first AUG in the construct, which we refer to as the canonical start, as opposed to cryptic downstream translation initiation. To do this, we compared WT reporter constructs to WT-STOP mutants, in which the canonical start was followed by two in-frame stop codons (Fig 1A). These downstream stop codons substantially reduced reporter expression in all three systems, suggesting that most translation initiated either at, or upstream of, the canonical start (Fig 1B).

**Figure 1:**
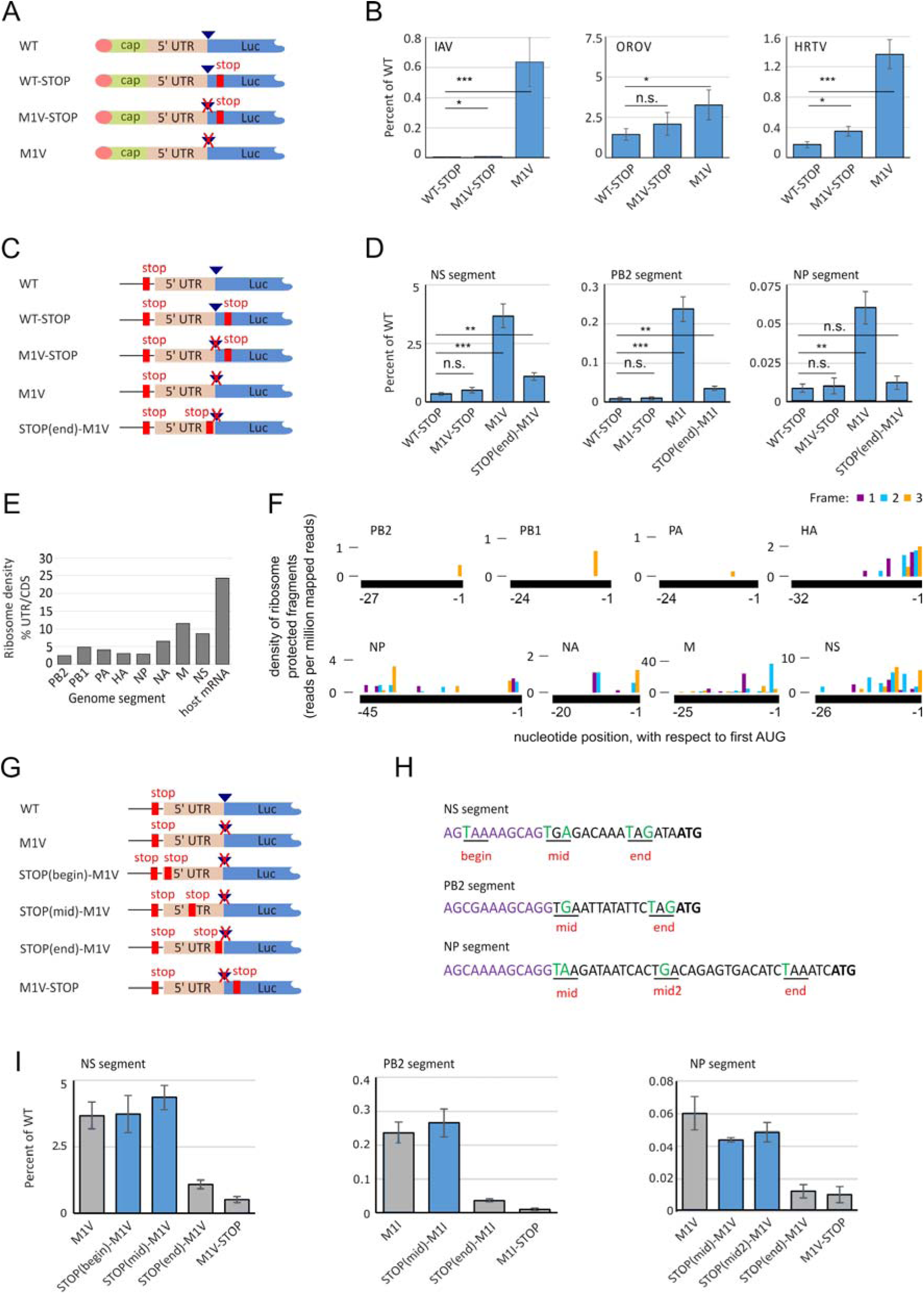
Upstream Translation Initiation in the 5′ Untranslated Regions of Segmented Negative-Strand RNA Viruses. (A) Schematic showing (in coding sense) the 5′ termini of viral reporter RNAs, in which a viral untranslated region (UTR) flanks a luciferase (Luc) reporter gene. Reporter RNAs were used to assess upstream translation in the mRNAs of influenza A virus (IAV), Heartland virus (HRTV) and Oropouche virus (OROV). The 5′ terminus of the mRNAs consisted of cap-snatched sequence from host mRNAs (cap), followed by a viral 5′ UTR (5′ UTR) and the reporter gene (Luc). Cap structures are indicated as circles, the most N-terminal AUG as a triangle, AUG mutations as crosses and stop codons as lines. (B) Luc expression when these reporters were included in minireplicon assays, as a percentage of expression with the WT construct, showing the means and s.d. of 5 (IAV), 3 (HRTV) or 4 (OROV) repeats compared to WT-STOP by Student’s 2-tailed t-test (n.s.: p ≥ 0.05, * p < 0.05, *** p ≤ 0.0005). (C) 5′ viral UTRs and Luc reporter genes were inserted, in coding sense, into a cellular RNA Polymerase II (RNAPII) transcribed plasmid in order to assess upstream translation of IAV segments 1, 5 and 8. A schematic showing mutations in the UTR region is shown. (D) Luc expression when these reporters were transcribed by RNAPII, as a percentage of WT. Means and s.d. of 3 repeats; compared to WT-STOP by Student’s 2-tailed t-test. (n.s.: p ≥ 0.05, * p < 0.05, ** p < 0.005, *** p ≤ 0.0005). (E) Summary Ribo-Seq measurements of viral and host mRNA from IAV infected cells, showing the ribosome density on 5′ UTRs as a percentage of its density on the main coding sequence (CDS). (F) Ribo-Seq profiles of the virally-encoded 5′ UTRs (black lines), with histograms showing the frequency of ribosomal P-site positions, inferred from the density of ribosome-protected fragments. Reads in frame with, +1 to, or +2 to the main ORF are shown in purple, blue and yellow, respectively. Ribo-Seq profiles of the canonical CDSs are shown in Supplementary Figure S1. (G) Schematic showing expression constructs, as in (C), modified to include additional stop codons to map upstream translation in IAV segments 1, 5 and 8. (H) 5′ UTR sequences of IAV segments 1, 5 and 8 showing the conserved sequence (purple), segment-specific sequences (black) and stop mutations (green) and stop codons (underlined). (I) Luc expression when these reporters were transcribed by cellular RNAPII, as a percentage of WT. The means and s.d. of 3 repeats are shown, with grey bars indicating data also included in panel (D).

We next compared translation initiation at the canonical start with initiation upstream of this position. To do this, we suppressed translation from the canonical start by mutating it from AUG to GUA (M1V). As expected, when followed by stop codons (M1V-STOP) these mutants behaved similarly to WT-STOP, with only a slight decrease in reporter attenuation (Fig 1B). However, in the absence of downstream stop codons, M1V mutants gave substantially higher reporter expression than WT-STOP (Fig 1B). For OROV and HRTV this increase was 2.3- and 8.0-fold, respectively, while for IAV the increase was 125-fold (Fig 1B). This showed that appreciable levels of translation can initiate from the 5′ ends of sNSV transcripts, even in the absence of a canonical start codon.

This unexpected reporter expression could be explained in two ways: by upstream translation initiation, or by non-canonical initiation from the mutated start codon. Distinguishing between these possibilities required us to mutate the viral 5′ UTR, but doing so in a minireplicon system is problematic as UTR sequences are required for normal viral transcription. To remove the need for processing by viral transcriptional machinery, we took the reporter construct with the most pronounced M1V phenotype, IAV NS, and cloned it in the positive sense into the cellular RNA polymerase II-transcribed mRNA expression vector pcDNA3A [31], introducing two in-frame stop codons upstream of the cloning site to exclude translation initiation from vector sequences (Fig 1C). Transfection of this construct into cells resulted in robust reporter expression. As in the minireplicon system, the WT-STOP and M1V-STOP mutations strongly attenuated reporter expression and the attenuation of M1V was partially alleviated when the downstream stop was removed (Fig 1D). The same effect was observed with the 5′ UTRs of the IAV PB2 and NP segments (segments 1 and 5, respectively; note that for segment 1 the start codon was abolished with an AUG to AUC mutation (M1I) to avoid creating a new AUG; Fig 1D). This suggested that translation initiation within a viral 5′ UTR was a property of the sequence itself and did not necessarily require some special property of the viral transcriptional machinery.

As we could now mutate sequences in the viral 5′ UTR without compromising transcription, we introduced in-frame stop codon towards the 3′ end of the 5′ UTRs, just upstream of the mutated start sites (STOP(end)-M1V; Fig 1C). For each of the three segments tested, expression from STOP(end)-M1V was substantially less than expression from M1V, suggesting that the increased reporter expression of M1V constructs compared to WT-STOP constructs was largely attributable to translation initiation upstream of the mutated start codon rather than from the mutated start codon itself (Fig 1D). We concluded that translation initiates at appreciable levels within the 5′ UTR sequences of IAV genes.

We next wished to look for evidence of upstream translation initiation in actual viral infections. To do this we infected cells with IAV, immobilised translating ribosomes by flash freezing, and performed ribosomal profiling (Ribo-Seq) of mRNAs. We were able to map assembled ribosomes to the UTRs of mRNAs transcribed from all eight viral segments, as well as to the UTRs of host mRNAs (Fig 1E, F; Supplementary Fig S1). Our measures of ribosome density are only semi-quantitative, but given that only one third of positions in the UTR would be in-frame with a reporter gene, the ratios of ribosomal density on UTRs to that on the canonical gene is consistent with the levels of upstream reporter translation observed in our minireplicon assays. We concluded that upstream translation initiation can occur in the 5′ UTRs of sNSV mRNAs.

### Upstream translation initiates from segment-specific viral untranslated regions

We next mapped sites of upstream translation initiation. Using IAV pcDNA3A reporters, we compared the effects of in-frame stop codons at different positions in the 5′ UTR of the M1V constructs – at the 5′ end of the viral UTR sequence (STOP(begin)-M1V); at a point between the terminal 12 nt, which are conserved across all IAV segments, and the segment-specific UTR sequence downstream of this (STOP(mid)-M1V); and at a point towards the 3′ end of the viral 5′ UTR (STOP(end)-M1V; Fig 1G,H). For all three segments tested, stop codons at the beginning and middle of the sequence did not reduce reporter expression, indicating that translation initiation in the viral 5′ UTRs occurred mainly in the segment-specific 3′ end of the UTR, rather than in the conserved 5′ end (Fig 1I).

Comparing reporter expression from the STOP(mid)-M1V and STOP(end)-M1V constructs suggested that the efficiency of upstream translation initiation in these segment-specific sequences varied between segments, with the luciferase signal attributable to the (mid)-(end) regions of segments 8, 1 and 5 (NS, PB2 and NP) in the ratio 115 : 15 : 1 (Fig 1I). As a further test for sequence specificity, we inserted a stop codon in the middle of the long segment-specific UTR of IAV segment 5 (NP; STOP(mid2)-M1V). This had little effect on reporter expression indicating that, for this segment, translation of the 5′ UTR initiated mainly towards the 3′ end of the 5′ UTR. We therefore concluded that when translation initiates in the 5′ UTRs of IAV mRNAs it does so in a segment-specific, sequence-dependent manner.

### Additional upstream translation initiates through ‘start-snatching’

We next sought to map upstream translation initiation within viral transcripts. When we examined the precise location of ribosomes in viral UTRs using our Ribo-Seq data, we found that the strongest ribosomal density was typically at the 3′ ends of the UTRs, consistent with the results of our reporter assays (Fig 1F). However, for the segments that returned the highest number of ribosome-protected fragments (RPFs; segments 5, 7 and 8, encoding the NP, M and NS genes respectively) it was possible to detect ribosomes even in the conserved terminal 5′ UTR sequences. This could not be explained by translation initiation in the segment-specific region of the 5′ UTR, as observed in our reporter assays, and suggested that in the context of viral infection there was an additional mechanism for translation initiation even further upstream.

We searched for further evidence of upstream translation during IAV infection by looking for evidence of novel viral proteins. To do this we purified IAV virions, extracted proteins, digested these with trypsin and searched for evidence of N-terminally extended viral proteins by mass spectrometry, broadening our search parameters to include proteins initiating at non-AUG positions. We repeatedly identified peptides that mapped to an in-frame translation of the UTR of segment 5 (NP), giving an N-terminally extended protein we termed NP-UTR (Fig 2A, B; Supplementary Fig S2, Supplementary Table S1). The peptides mapping to the UTR sequence were detected at only a small proportion of the average intensity of peptides mapping to the downstream NP sequence, suggesting that NP-UTR was a low-frequency variant of NP (Fig 2C). Notably, peptides identifying NP-UTR spanned almost the full extent of the NP 5′ UTR, including the conserved 5′ sequences (Fig 2B). Indeed, as the N-termini of these tryptic peptides mapped to tryptic cleavage sites, it is probable that their translation initiated still further upstream. This, combined with the lack of translation initiation within the terminal 5′ sequence observed in minireplicon assays, suggested that translation of IAV mRNAs could initiate at sites upstream of the viral 5′ UTR.

**Figure 2:**
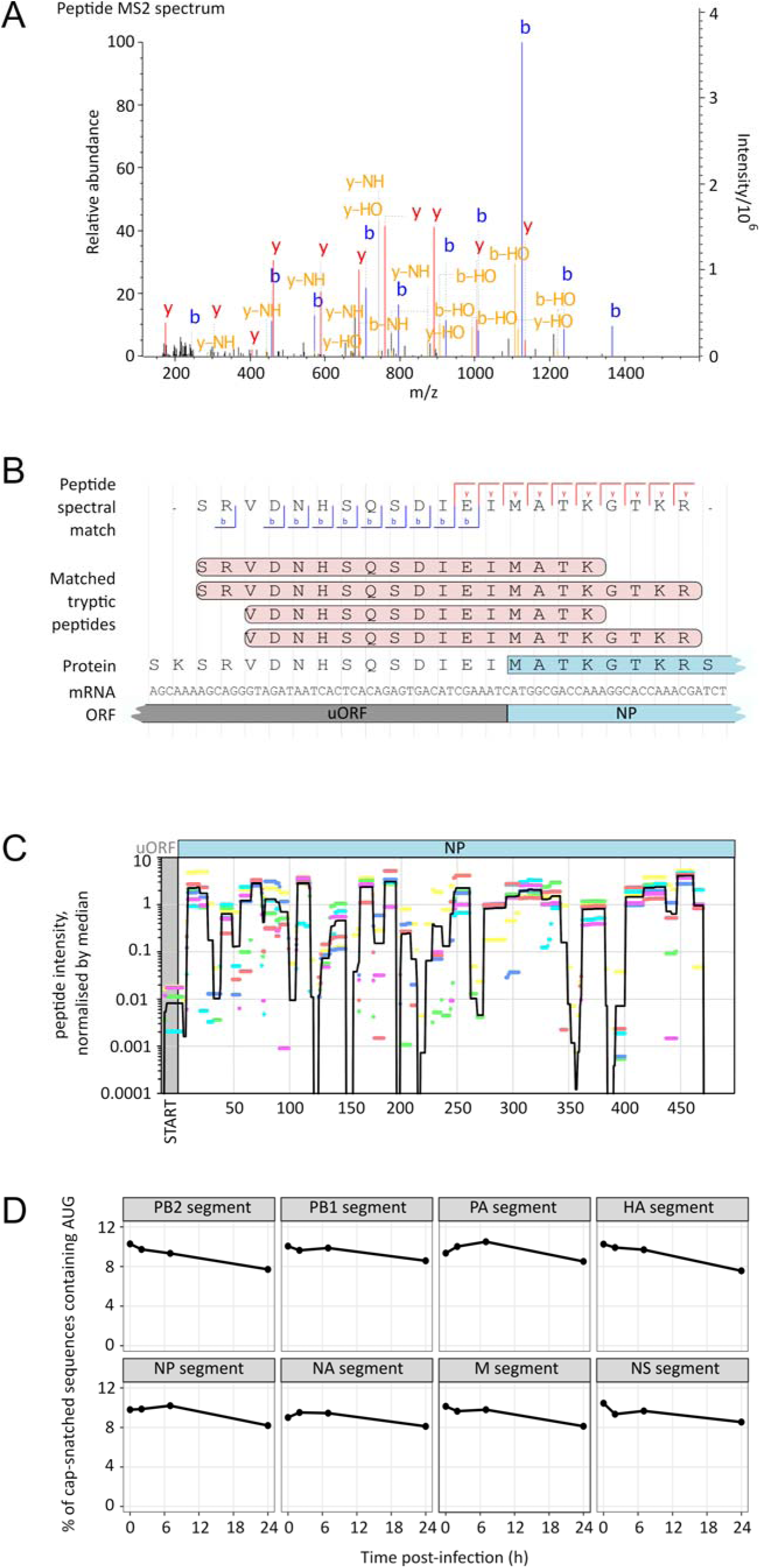
Upstream Translation Initiation of Virally-Encoded UTRs. (A) – (C) IAV particles were purified and subjected to tryptic digest, liquid chromatography and tandem mass spectrometry. (A) A representative fragment mass spectrum (MS2) describing a tryptic peptide mapping to the 5′ UTR of IAV segment 5 (NP). (B) Interpretation of this mass spectrum, comparison of its sequence to other, overlapping tryptic peptides that were identified (see Supplementary Fig S2), and comparison of these peptides to the NP gene and 5′ UTR of IAV segment 5. (C) The intensities (MS1) with which peptides were detected at positions spanning the entirety of IAV segment 5. Data from 6 separate experiments (colour-coded) are shown, normalised to median intensity of peptides mapped to NP and its 5′ UTR. The average intensity mapped to each residue is indicated with a black line and the region spanning the upstream ORF (uORF) is shaded. (D) Cap Analysis Gene Expression (CAGE) sequencing of cap-snatched sequences for the eight segments of the IAV genome, collected from infected cells at various times post-infection and showing the percentage of cap-snatched sequences containing AUG codons.

The apparently contradictory finding of translation initiation upstream of the 5′ UTR is plausible for an sNSV, as these viruses produce mRNAs by cap-snatching. We reasoned that start codons within the cap-snatched leader could provide additional sites for upstream translation initiation. These sites would be appended to mRNAs produced by the viral transcriptional machinery, but would be absent in the products of our pcDNA3A reporter assay which were transcribed by cellular RNA Polymerase II. We therefore asked whether IAV cap-snatched leader sequences contained AUG codons at a high enough frequency to explain the translation of proteins such as NP-UTR. To do this, we infected cells with IAV and sequenced viral mRNAs by Cap Analysis Gene Expression (CAGE) sequencing. Because IAV infections profoundly alter the mRNA pool of IAV infected cells [22, 32], we sequenced mRNA at timepoints throughout the first 24 h of infection. We found that approximately 10% of cap-snatched leader sequences contained an AUG (Fig 2D; Supplementary Fig S3). This proportion was consistent for mRNAS from all IAV genome segments and declined only slightly over the first 24 h of infection (Fig 2D) without an obvious preference for AUG position within the leader (Supplementary Fig S4). We then compared our results to a previously-published deep-sequencing description of IAV cap-snatched sequences [20]. Despite differences in methodology, we were able to identify AUGs in a comparable percentage of the cap-snatched leaders of viral mRNAs when using data from this study (Supplementary Fig S5).

Thus, cap-snatching by IAV can acquire start codons suitable for the upstream translation of the entire virally-encoded UTR, a process which would account for our detection of the extended NP-UTR protein in virions. We termed this additional upstream translation initiation process ‘start-snatching.’

### Upstream translation initiation allows the expression of cryptic viral open reading frames

When we considered the two modes of upstream translation initiation that we had identified in sNSVs, we realised that both of them could create novel viral gene products in two different ways.

One would be to create short N-terminal extensions to a segment’s major gene product(s) by translating the 5′ UTR, as we had observed with IAV NP-UTR. Sequence analysis of the strains used in our minireplicon assays showed that this would be possible for six of the eight genome segments of the IAV strain used (influenza A/WSN/33(H1N1); WSN) and two of the three segments of OROV and HRTV; in the other segments the 5′ UTR contained an in-frame stop codon.

The other route for creating novel gene products would be if translation initiated out of frame with the major gene product, accessing a cryptic overlapping 5′ reading frame (Fig 3A; Supplementary Fig S6). In the strains used in our minireplicon assays, we found overlapping 5′ ORFs encoding polypeptides that were comparable in length to previously identified short proteins (defined here as > 20 codons [33–35]) in frame 3 of IAV segments 1 – 4 (PB2, PB1, PA and HA) and of OROV segments 1 – 3 (L, M and S). In contrast, the overlapping 5′ ORFs in HRTV were shorter. Because of this difference between OROV and HRTV we searched for overlapping 5′ ORFs in the complete genome sequences of 78 different peribunyavirus species, and we found that there were overlapping 5′ ORFs of >20 codons in 62% of L genes, 90% of M genes and 73% of S genes (Supplementary Fig S7, Supplementary Tables S2 – S4). In the case of IAV considerably more sequence data were available, allowing us to determine the conservation of the overlapping 5′ ORFs in frame 3. To do this we compared between 23,000 and 43,000 IAV strains, depending on the segment. We found that the majority of frame 3 ORFs for each segment were of consistent lengths, and that the sequences they encoded were highly conserved (Fig 3B). We concluded that sNSVs often have the genetic potential to access cryptic 5′ uORFs through upstream translation initiation, and that in the case of IAV these uORFs can be highly conserved.

**Figure 3:**
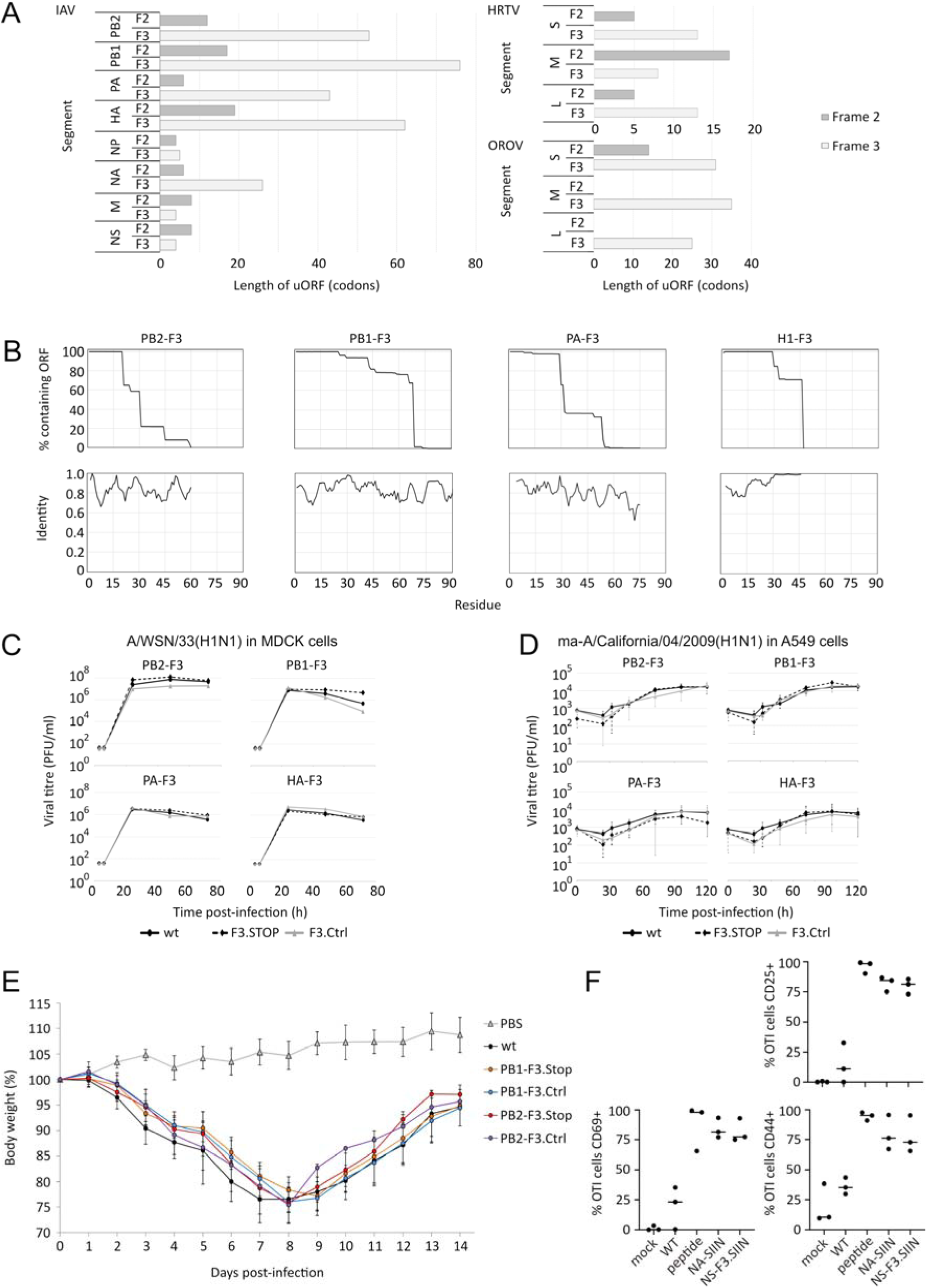
Upstream Translation Initiation Can Access Overlapping Viral ORFs. (A) The length of 5′ ORFs in IAV (A/WSN/33; WSN), OROV and HRTV, beginning at the extreme 5′ end of each segment (coding sense, including the 5′ UTR) and continuing in frames 2 and 3 until encountering a stop codon. (B) Comparison of 5′ frame 3 (F3) sequences from segments 1 – 4 of >20,000 IAV isolates showing the proportion that do not encounter a stop codon by a given length (upper) and the degree of identity of the encoded sequences (lower). (C – D) Viruses containing F3.STOP or synonymous F3.Ctrl mutations within the PB2, PB1, PA and HA segments were created by reverse genetics. To assess multi-cycle growth kinetics, media were collected from cells infected at an initial MOI of 0.01 and plaque titres determined on MDCK cells. Growth kinetics were determined when (C) mutants of the laboratory-adapted IAV WSN were used to infect MDCK cells and (D) mutants of the near-clinical IAV isolate mouse adapted-A/California/04/2009(H1N1) (maCa04) were used to infect human lung A549 cells (means and s.d. of 3 experiments). (E) Selected maCa04 viruses were used to infect BALB/c mice and mouse weight was recorded over time. (F) IAV antigen preparations were obtained by infecting MDCK cells with WT virus, virus with a SIIN peptide in the stem of the NA protein (NA.SIIN) or virus with a cryptic SIIN peptide in F3 of segment 8 (NS-F3.SIIN). Bone marrow derived dendritic cells (BMDCs) were treated overnight with the IAV antigen preparations and then co-cultured with SIIN-specific OTI T cells. OTI activation was determined by immunostaining for the T cell activation markers CD25+, CD44^HIGH^+ and CD69+. Mock infection and exposure of BMDCs to a purified SIIN peptide were used as controls. Datapoints and means from 3 experiments are shown.

We wished to determine whether the conserved overlapping frame 3 (F3) ORFs in segments 1 – 4 of IAV, which we termed PB2-F3, PB1-F3, PA-F3 and HA-F3, respectively, were critical for viral replication. We tested this by reverse genetics, creating viruses with point mutations near the start of the F3 ORFs that were synonymous in frame 1, and which in frame 3 could be either conservative mutations (F3-Ctrl) or stop mutations (F3-STOP; Supplementary Fig S8A). As WSN is a heavily laboratory-adapted strain of the virus we also mutated the mouse-adapted near-clinical isolate, influenza virus A/California/04/2009(H1N1) (maCa04; Supplementary Fig S8B). When mutating frame 3, we observed no differences between the growth kinetics of WT, F3-STOP and F3-Ctrl viruses, either when growing WSN in MDCK cells (Fig 3C) or when growing maCa04 in human lung A549 cells (Fig 3D). We also infected mice with the mutant PB2-F3 and PB1-F3 maCa04 viruses, and observed no differences in pathogenicity between the WT, F3-STOP and F3-Ctrl viruses (Fig 3E). Thus, although the coding potential of F3 proteins is highly conserved in naturally circulating strains of influenza virus (Fig 3B), they appear to be dispensable in laboratory conditions. We note that this, like their apparently low levels of expression, is consistent with the properties of previously-characterised IAV accessory proteins [6].

In addition to modulating viral gene expression directly, viral proteins within infected cells are presented to the immune system by MHC I. If conserved, as the IAV F3 proteins appear to be, this can provide targets for immune responses in future infections. To ask whether F3 proteins could be presented by MHC I and recognised by the immune system, as well as to test whether in principle F3 ORFs could be translated, we created a modified IAV containing a frame 3 insertion of OVAI (OVA 257-264; SL8; SIINFEKL), a class I-restricted epitope of ovalbumin. We inserted OVAI into segment 8 (NS) of the virus, deleting a number of naturally-occurring F3 stops to create a cryptic 5′ ORF so that we could exploit the tolerance of a linker region in the NS1 gene to insertion mutations (NS-F3.SIIN; Supplementary Fig S9) [36]. To ask whether the F3 5′ ORF was translated we treated bone marrow derived dendritic cells (BMDCs) with a sonicated IAV antigen preparation and then incubated these with purified CD8 T cells from OTI mice (Fig 3F). Little or no OTI activation was observed with mock-infected DCs, but the NS-F3.SIIN virus preparation resulted in robust OTI activation, as measured by levels of the activation markers CD44, CD25 and CD69. This was comparable to the activation seen following treatment with purified OVAI peptide or to a control experiment using a virus in which we had inserted OVAI, in frame 1, into the stem region of the viral NA protein (NA-SIIN; [37]). We concluded from this that cryptic 5′ ORFs of sNSVs, accessible through upstream translation initiation, can be expressed by infected cells, and moreover that the products of these cryptic ORFs can be presented to the immune system.

## Discussion

Our results show that upstream translation initiation, a process that regulates the expression of a large proportion of cellular genes [10,12–15,38], is also exploited by the sNSVs that parasitize those cells. This finding is of general relevance to virology: given our finding that translation can initiate even in the short virally-encoded 5′ UTRs of IAV (Fig 1 G-I), it is plausible that upstream initiation occurs in the 5′ UTRs of many other eukaryotic viruses. Indeed, it is important to note that the viral UTRs we examined in this study do not contain AUG codons, meaning that translation initiation in these regions is presumably from non-AUG codons, a common if low-level effect in both host and viral genes [4,39,40]. Our data therefore add to a growing number of reports of upstream translation initiating on virally-encoded sequences, in some cases with clear regulatory effects [4,41–46]. On the basis of our results we would argue that upstream translation of viral genes should not be discounted, even in the absence of virally-encoded AUGs.

The reliance of sNSVs on cap-snatching for mRNA synthesis makes upstream translation initiation in these viruses particularly interesting. Our data indicate that for sNSVs, upstream translation can initiate not only on virally encoded sequences, but also in the short host-encoded sequences appropriated by cap-snatching (Fig 2). This ‘start-snatching’ would require functional start codons extremely close to the 5′ end of cellular mRNAs. Start codons of this sort, referred to as Translation Initiator of Short 5′ UTR (TISU) motifs, have been reported in approximately 4 % of mammalian mRNAs, being particularly common in ‘housekeeping’ genes [16–18]. We note that this figure is comparable to the ribosomal density we observed on the 5′ UTRs of viral mRNAs (Fig 1E) and is compatible with the 8 – 10 % of viral cap-snatched leaders which we found to contain AUGs (Fig 2D). TISU motifs are functional within 30 nt of the 5′ cap, including at sites at the extreme end of mRNA with 5’ UTRs as short as 5 nt [16–18]. As with other upstream translation sites, TISUs have been proposed to regulate the expression of downstream genes, in response to factors including energy deprivation and the circadian rhythm [47–49]. As a result sNSVs, being capable of start-snatching, will necessarily incorporate host regulatory elements into their own mRNAs. We propose that, in this way, start-snatching may allow sNSVs to modulate their translational profile in response to the effects that infection is having on their host’s gene expression.

Our data also show that upstream translation in sNSVs can lead to the expression of novel viral proteins. We have shown this experimentally for IAV (Fig 2A-C, Fig 3F) and it is clear from genome sequences that the genetic potential to generate proteins from upstream translation is widespread in IAVs and peribunyaviruses (Fig 3A, B, Supplementary Fig S6 and Supplementary Tables S3, S4). Upstream translation could lead to N-terminal extensions to canonical proteins, as for the NP-UTR protein we detected in IAV virions (Fig 2 A-C). It could also lead to the translation of proteins from cryptic overlapping uORFs, which have the potential to be presented to the adaptive immune system (Fig 3F). While this work was being completed we became aware of another study, currently in pre-print form, that examined translation initiation from the cap-snatched sequences of IAV [50]. Consistent with our results, the authors of this work also report the expression of short uORFs from a number of IAV genome segments, producing proteins they referred to as Upstream Flu ORFs (UFOs). Novel viral proteins of this sort share several key features. They would most likely be expressed at low to moderate levels – certainly lower than most viral proteins, although the proteins of rapidly-replicating viruses such as IAV are typically expressed at extremely high levels. Based on our data, this appears to be the case for NP-UTR (Fig 2C). If they overlap a canonical ORF they will also be short, typically less than 100 codons (Fig 3A, Supplementary Fig S6). This is shorter than most viral and cellular proteins, though comparable in length to many short functional ORFs in viral and cellular genomes [33–35,38,51,52]. Finally and, given their small size and moderate expression levels, perhaps surprisingly, the overlapping uORFs in segments 1 – 4 of IAV are highly conserved.

The conservation of uORFs makes it tempting to suggest that they might encode functional proteins. However, arguments of this sort must be treated with caution, as other forms of selection also act on IAV genome sequences. In particular, genome packaging signals in the primary RNA sequence are concentrated in the terminal regions of each segment [53–55], resulting in a suppression of synonymous codon usage comparable to that of known overlapping ORFs (Supplementary Fig S10) [54, 56]. One can instead look for suppression of stop codons in the +1 and +2 frames (reasoning that conservation of primary RNA sequence should not discriminate specifically against stop codons), but this requires some care as the occurrence of stop codons in these frames can be affected by codon usage in the main ORF, as well as by nucleotide and dinucleotide biases [3,57,58]. We attempted to address this by using randomization to assess the likelihood that the absence of stop codons in any given region could have arisen through chance alone (Supplementary Fig S10; see Methods for details). Of the areas where stop codons appeared to be suppressed, the only ones that could not plausibly be explained by chance alone were the known +1 frame M2 and NEP ORFs and a short +2 frame region near the 5′ end of the NS1 ORF (Supplementary Fig S10). Clearly this is a fairly weak analysis, best suited to identifying long ORFs conserved between highly divergent sequences. Notably it did not score as significant a number of established overlapping ORFs, such as the PB1-F2 and PA-X [59, 60]. However, this analysis and the known selection for primary RNA sequence in IAV genome segments mean that the observed conservation of uORFs is uninformative regarding their functional importance. As a result, while it is clear that uORFs can be translated, whether there is direct fitness benefit arising from the proteins they encode must be assessed experimentally on a case-by-case basis.

Whether or not the virus makes direct use of uORF proteins, it is clear that the adaptive immune system can recognise epitopes from within their reading frames (Fig 3F). MHC I presentation of uORF-derived peptides poses the risk of an adaptive immune response developing against sNSVs, analogous to the risks posed to IAV by the presentation of alternative reading frames (ARFs) and defective ribosomal products (DRiPs) [61–65]. Indeed, the risks posed by the presentation of uORFs are potentially even higher due to the high conservation of these sequences. They are conserved nonetheless, which suggests that any cost the virus incurs through their visibility to the immune system is outweighed by the fitness benefits of maintaining this genetic architecture.

In summary, we have shown that sNSVs can expand their genetic repertoire through a widespread capability for upstream translation initiation. Like all viruses, sNSVs rely on their hosts for the translation of their genes, and by utilising their host’s mechanisms of upstream translation this diverse group of viruses have gained the ability to develop additional layers of gene regulation and to potentially encode further gene products in their highly constrained, short RNA genomes.

## Materials and Methods

### Cells and viruses

Madin–Darby Canine Kidney (MDCK) cells, A549 human lung epithelial cells, and 293T human embryonic kidney cells were cultured in Dulbecco’s Modified Eagle’s Medium (DMEM; Gibco) supplemented with 10% foetal bovine serum (FBS; Gibco). Madin-Darby Bovine Kidney (MDBK) cells were cultured in Minimum Essential Medium (MEM; Sigma) supplemented with 2 mM L-glutamine and 10% foetal calf serum (FCS). BSR-T7/5 golden hamster cells were cultured in Glasgow Minimal Essential Medium (GMEM) supplemented with 10% FCS and 10% tryptose phosphate broth under G418 selection. All cells were maintained at 37 C and 5% CO_2_.

The influenza A viruses A/Puerto Rico/8/34(H1N1) (PR8) and mouse-adapted A/California/04/09(H1N1) (maCa04) were generated by reverse genetics and propagated on MDCK cells, as described previously [66]. The influenza virus A/WSN/33(H1N1) (WSN)was propagated on MDBK cells and the influenza virus A/Udorn/72(H3N2) (Udorn) was propagated on MDCK cells, as described previously [22, 67]. IAVs, with the exception of WSN, were propagated in the presence of 1 µg/ml TPCK-trypsin. Plaque assays were carried out in MDCK cells and visualised by immunocytochemistry, as previously described [68].

### Plasmids

Plasmids used for IAV minireplicon assays were the firefly-luciferase-encoding phPOLI-NS-Luc [69]; the viral-gene-encoding pcDNA-PB1, pcDNA-PB2, pcDNA-PA, pcDNA-NP (a kind gift of Prof Ervin Fodor, University of Oxford) [31]; the Renilla-luciferase-encoding control plasmid pRL-TK (Promega) and empty vector pcDNA3A. Plasmids used for HRTV minireplicon assays were the Renilla-luciferase-encoding pT7HRTMRen(–); the viral-gene-encoding pTMHRTN and pTMHRTL and the firefly-luciferase-encoding control plasmid pTM1-FFluc [70]. Plasmids used for OROV minireplicon assays were the Renilla-luciferase-encoding pTVT7-OROMhRen(–); the viral-gene-encoding pTM1-ORO-N and pTM1-ORO-L; the firefly-luciferase-encoding control plasmid pTM1-FFluc and the empty vector pTM1 [71]. For pcDNA IAV reporter assays, the NS-Luc sequence from phPOLI-NS-Luc was cloned into pcDNA to create pcDNA-NS-Luc. The 5′ UTR sequence of this plasmid was edited by site directed mutagenesis to produce pcDNA-PB2-Luc and pcDNA-NP-Luc. Plasmids used for reverse genetics were the PR8 pDUAL plasmids (a kind gift of Prof Ron Fouchier, Erasmus MC) [72] and the maCa04 pDP2002 plasmids (a kind gift of Prof Daniel R. Perez (University of Georgia, USA) [73]. Site-directed mutagenesis of plasmids was performed using the Q5 site-directed mutagenesis kit (Qiagen); the edited NS segment sequence required for the PR8-NS.F3.SIIN mutant virus (described in Supplementary Fig S9A) synthesised by Genewiz.

### Minireplicon assays and luciferase assays

Minireplicon assays were performed as previously described [70,71,74]. Briefly, and using the plasmids indicated above, for IAV Lipofectamine 2000 (Invitrogen) was used to transfect sub-confluent 293T cells, and for HRTV and OROV LT-1 transfection reagent (Mirus) was used to transfect sub-confluent BSR-T7/5 cells. To measure luciferase expression from pcDNA constructs, Lipofectamine 2000 (Invitrogen) was used to transfect sub-confluent 293T cells with the plasmids indicated above. In all cases, after 24 h cells were processed using a Dual-Luciferase Reporter Assay System (Promega), with luciferase measured using Glowmax 20/20 luminometer (Promega).

### Ribo-Seq analysis

A549 cells were grown on 100-mm dishes to 90% confluency and infected with PR8 at a multiplicity of infection (MOI) of 5. At 5 h p.i., cells were rinsed with 5 ml of ice-cold PBS, flash frozen in a dry ice/ethanol bath and lysed with 400 μl of lysis buffer [20 mM Tris-HCl pH 7.5, 150 mM NaCl, 5 mM MgCl_2_, 1 mM DTT, 1% Triton X-100, 100 μg/ml cycloheximide and 25 U/ml TURBO DNase (Life Technologies)]. The cells were scraped extensively to ensure lysis, collected and triturated ten times with a 26-G needle. Cell lysates were clarified by centrifugation at 13,000 g for 20 min at 4°C. Lysates were subjected to Ribo-Seq based on previously reported protocols [46,75,76]. Ribosomal RNA was removed using Ribo-Zero Gold rRNA removal kit (Illumina) and library amplicons were constructed using a small RNA cloning strategy adapted to Illumina smallRNA v2 to allow multiplexing. Amplicon libraries were deep sequenced using an Illumina NextSeq500 platform (Department of Biochemistry, University of Cambridge). Ribo-Seq sequencing data have been deposited in ArrayExpress (http://www.ebi.ac.uk/arrayexpress) under the accession number E-MTAB-8405.

For the Ribo-Seq computational analysis, adaptor sequences were trimmed from Ribo-Seq reads using the FASTX-Toolkit (hannonlab.cshl.edu/fastx_toolkit/) and reads shorter than 25 nt following adaptor trimming were discarded. Trimmed reads were mapped sequentially the genomes of the (*Homo sapiens*) and virus (PR8, with accession numbers EF467817, EF467818, EF467819, EF467820, EF467821, EF467822, EF467823 and EF467824) using bowtie version 1 [8], with parameters -v 2 --best (i.e. maximum 2 mismatches, report best match). Mapping was performed in the following order: host rRNA, virus RNA, host RefSeq mRNA, host non-coding RNA and host genome. The host databases used were rRNA: NR_003287.2, NR_023379.1, NR_003285.2, NR_003286.2; mRNA: 35809 human mRNA National Center for Biotechnology Information (NCBI) RefSeqs, downloaded 24 Jan 2013; ncRNA: Ensembl Homo_sapiens.GRCh37.64.ncrna.fa; genome: UCSC hg19.

To account for different library sizes, reads per million mapped reads (RPM) values were calculated using the sum of both positive-sense host mRNA reads and virus RNA reads as the denominator. For viral genome coverage plots, and for meta-analyses of host mRNA coverage, mapping positions were obtained from the 5′ end of RPFs plus a 12-nt offset, so as to approximate the location of the ribosomal P-site. Length distributions of RPFs were obtained from reads that mapped entirely within host coding regions. Histograms of RPF positions relative to host mRNA initiation and termination codons were derived from reads mapping to mRNAs with annotated coding regions ≥450 nt in length and with annotated UTRs ≥60 nt in length (Supplementary Fig S1B). These RPFs were also used to determine the relative read densities in host 5′UTRs and coding sequences; in this case only reads with estimated P-sites mapping within the 3′-most 60 nt of the 5′ UTR and the 5′-most 450 nt of the coding region were counted.

To quantify RPFs that spanned the junction between virus-derived and host-derived (i.e. cap-snatched fragment) sequence, reads containing the conserved motif GC[AG]AAAGCA (i.e. nt 2 – 10 of virus genome segments), and with at least 7 nt of sequence 3′ of this motif, were identified within the set of previously unmapped reads. If the 5′ end of this 3′ region could be mapped unambiguously to a virus segment (from nt 11 onwards), then the read was considered to have been derived from that segment. This enabled extension of the RPF estimated-P-site density plots on virus mRNAs as far 5′ as nt 2 of each segment, since a 28-nt RPF with P-site mapping to nt 2 – 4 would normally have a 3′ end mapping to approximately nt 17.

### Mass spectrometry

The purification of influenza virions and collection of mass spectra by LC-MS/MS has been described previously [67], and followed previously-described protocols for purification, mass spectrometry and data analysis [77]. Briefly, the IAV WSN was propagated on MDBK cells. Six viral stocks were prepared, of which half were subjected to haemadsorption on chicken red blood cells to stringently remove non-viral material. Virus particles were then purified by sucrose gradient ultracentrifugation, lysed in urea, reduced, alkylated and digested with trypsin and LysC. Tryptic peptides were analysed by liquid chromatography and tandem mass spectrometry (LC-MS/MS) using an Ultimate 3000 RSLCnano HPLC system (Dionex, Camberley, UK) run in direct injection mode and coupled to a Q Exactive mass spectrometer (Thermo Electron, Hemel Hempstead, UK) in ‘Top 10’ data-dependent acquisition mode. Raw files describing these mass spectra have been deposited at the Mass spectrometry Interactive Virtual Environment (MassIVE; Center for Computational Mass Spectrometry at University of California, San Diego) and can be accessed at http://massive.ucsd.edu/ProteoSAFe/datasets.jsp using the MassIVE ID MSV000078740. For the purposes of this project, data were re-analysed using MaxQuant 1.5.8.3 analysis software [78] using standard settings and the following parameters: label-free quantitation and the iBAQ algorithm [79] enabled; enzyme: trypsin/P; variable modifications: oxidation (M) and acetyl (Protein N-ter); and fixed modifications: carbamidomethyl (C); digestion mode: semi-specific free N-terminus. Peptide spectra were matched to custom databases containing the WSN proteome (including full-length translations of all six reading frames), an edited version of the *Bos taurus* proteome (UP000009136; retrieved from UniProt on 16/05/2017) in which all instances of the ubiquitin sequence had been deleted, and a single repeat of the ubiquitin protein sequence.

### Sequencing of cap-snatched leader sequences

The sequencing of cap-snatched leader sequences was described in detail in a recent pre-print [22]. Briefly, primary CD14+ human monocytes were isolated from 4 volunteer donors under ethical approval from Lothian Research Ethics Committee (11/AL/0168) and cultured in the presence of 100 ng/ml (104 U/ml) recombinant human colony-stimulating factor 1 (a gift from Chiron, USA) for 8 days to differentiate them into macrophages. Monocyte-derived macrophages were then infected with influenza (Udorn) at an MOI of 5, harvested at 0, 2, 7 and 24 hours post-infection (times defined as starting after a 1h adsorption step), and processed for RNA extraction using a miRNeasy Mini Kit (Qiagen). Cap analysis of gene expression (CAGE) was performed as part of the FANTOM5 project, following the procedure of [80]. Data were processed as in [81] using custom Python scripts available at https://github.com/baillielab/influenza_cage ’ATG analysis’. The datasets analysed during the current study are available in the Fantom5 repository, http://fantom.gsc.riken.jp/5/data/

### Influenza A virus frame 3 conservation analysis

Full-length sequences for influenza A virus gene segments 1 (PB2, 42311 sequences), 2 (PB1; 42303 sequences), 3 (PA; 42756 sequences) and 4 (HA; H1 subtype only; 23798 sequences) were downloaded from the NCBI on 30/08/2019. Nucleotide multiple sequence alignments (MSAs) were created and clipped 450 nt downstream of the first AUG codon. All sequences were translated and frame 3 (+2) sequences were extracted; each frame 3 sequence was clipped after its first stop codon. Clipped frame 3 protein sequences were aligned by MAFFT using default parameters [82], with spurious or poorly aligned reads removed using trimAl, using parameters -resoverlap 0.70 -seqoverlap 75. Codon usage tables were compiled using BioEdit.

### Peribunyavirus uORF analysis

Representative reference sequences were downloaded from the NCBI in September 2018 for all species in the family *Peribunyaviridae*, as defined by the International Committee for Taxonomy of Viruses (ICTV). All available segments for each species were downloaded and examined. For each segment, we recorded the start and stop coordinates of the major ORF. To examine the coding potential of these sequences, we used custom scripts to virtually translate reference sequences in all three frames and identify regions of uninterrupted coding sequence that begin upstream of the start codon of the major ORF, which are not preceded by an in-frame stop codon. All scripts and data used in this analysis are openly available in an online repository (https://giffordlabcvr.github.io/Peribunyaviridae-GLUE/).

### Mouse pathogenesis studies

Six-week-old female BALB/c mice were anaesthetised with isofluorane before intranasal inoculation with 1000 PFU of each influenza virus (n=5). The animals were monitored daily for body weight changes and survival for 14 days after virus challenge. For ethical reasons, mice presenting ≥25% body weight loss were humanely euthanized. All procedures that required the use of animals performed in Portugal were approved by the Instituto Gulbenkian de Ciência Ethics Committee and the Animal Welfare Body, as well as by the Portuguese Authority for Animal Health, Direção Geral de Alimentação e Veterinária (DGAV).

### OTI T cell activation assay

IAV antigen was propagated by infecting MDCK cells with IAV PR8 wild type, PR8 containing an NS segment with SIINFEKL inserted into frame 3 (PR8-NS.F3.SIIN) or PR8 containing an NA segment with SIINFEKL inserted into frame 1 (PR8-NA.SIIN [37]). The IAV antigen preparations were prepared as described [83, 84]. Briefly, MDCK cells were infected for 48 h with each IAV stain and then centrifuged, resuspended in 0.1 M glycine buffer containing 0.9% NaCl (pH 9.75), and shaken at 4°C for 20 min. Preparations were sonicated 4 times at 10 s intervals before centrifugation, and the supernatant stored at -80°C.

Bone marrow was taken from 10-14 week old naïve female C57BL/6 mice, purchased from Envigo (UK) and maintained at the University of Glasgow under standard animal husbandry conditions in accordance with UK home office regulations and approved by the local ethics committee. Bone marrow derived dendritic cells (BMDCs) were prepared as previously described [84] and incubated overnight with IAV antigen preparations. Control BMDCs were incubated with SIINFEKL peptide (Ovalbumin (257-264), chicken, Sigma-Aldrich) for 1 h at 37°C.

OTI mice [85] were bred in-house on a mixed genetic background. Animals were kept in dedicated barriered facilities, proactive in environmental enrichment under the EU Directive 2010 and Animal (Scientific Procedures) Act (UK Home Office licence number 70/8645) with ethical review approval (University of Glasgow). Animals were cared for by trained and licensed individuals and humanely sacrificed using Schedule 1 methods. Lymph nodes (LN) (inguinal, brachial, axillary and cervical) and spleen were obtained from OTI mice sacrificed at weeks 12-13. CD8 T cells were negatively selected from LN and spleen using EasySep™ Mouse CD8+ T Cell Isolation Kit (Stemcell technologies).

BMDCs that had been exposed to viral antigen were co-cultured with CD8+ OTI T cells for 24 h. Activated T cells were detected by immunostaining with antibodies against Va2-E450 (Thermo Fisher), Vb5-PE (M59-4 BD Biosciences), CD8-Alexaflor488 (53-6.7 Thermo Fisher), CD25-APC (PC61.3 Thermo Fisher), CD44-PerCpC5.5 (IM7 Thermo Fisher), and CD69-PerCy7 (H1.2F3 Thermo Fisher). Data were acquired with a BD Fortessa cell analyser and analysed by FlowJo (BD, version 10).

### Analysis of stop codon suppression in influenza A viruses

All influenza A virus nucleotide sequences were downloaded from the NCBI on 28 July 2019. Patent sequence records, sequences with NCBI keywords “UNVERIFIED”, “STANDARD_DRAFT” or “VIRUS_LOW_COVERAGE”, and sequences with any ambiguous nucleotide codes (e.g. “N”s) were removed. leaving 109,132 sequences. Sequences were sorted into the eight segments using tblastn [86] with PR8-strain peptide sequences as queries.

For the three-frame stop codon plots, a set of “representative” sequences for each segment was obtained using BLASTCLUST (a single-linkage BLAST-based clustering algorithm; [86]) to cluster closely related sequences. One representative sequence from each cluster was selected. A disadvantage of this approach is that defective sequences (from defective viruses or from poor quality sequencing or misassembly) tend to form their own clusters and so become over-represented in the set of “representative” sequences. To guard against such problems, we used a number of restrictive selection criteria (see below). In segments 7 and 8, we first inserted “NN” immediately 5′-adjacent to the splice acceptor site in all sequences to fuse the coding regions of M2 and M1, and of NEP and NS1. We then identified the longest AUG-to-stop-codon open reading frame (ORF) in every sequence. We found the modal ORF length for each segment and discarded all sequences where the ORF was not of the modal length. This resulted in the loss of ∼37%, ∼25% and ∼15% of segment 4, 5 and 6 sequences, respectively, and <2% of sequences for other segments. For each segment, we then constructed the consensus amino acid sequence for the longest ORF and discarded all sequences which did not have a gap-free longest ORF amino acid alignment to the consensus. In this step, pairwise sequence alignments were performed with MUSCLE [87]. This step removed ∼43% of segment 4 sequences and ∼40% of segment 6 sequences, but only 1 other sequence among the other segments. All remaining sequences had >73% longest-ORF amino acid identity to the respective consensus sequence.

For each segment, we then clustered the longest-ORF amino acid sequences with BLASTCLUST (parameters -p T, -L 1, -b T). We applied BLASTCLUST with different amino acid identity thresholds (-S parameter) starting from 99.9% and stepping downwards in decrements of 0.1% until the number of clusters was 50 or fewer. We then excluded all clusters with just a single sequence. For each segment, we then chose the lowest BLASTCLUST identity threshold that resulted in ≥50 non-singleton clusters. This resulted in amino acid identity thresholds of 98.9%, 99.1%, 98.9%, 98.8%, 99.2%, 96.4%, 98.8% and 97.9%, and 58, 51, 50, 57, 64, 52, 56 and 56 non-singleton clusters, for segments 1–8 respectively. To choose a representative sequence for each cluster, we first extracted the nucleotide sequence corresponding to the longest ORF. To mitigate the effect of potential sequencing errors, in each cluster the representative sequence was chosen to be the sequence with the most identical copies (with ties broken arbitrarily) or, if there were no duplicated sequences, the sequence closest to the centroid (the minimum summed pairwise nucleotide distances from sequence *i* to all other sequences *j* within the cluster).

Since all selected coding-region sequences for each segment were of the same length, sequence alignment at this stage was trivial. Synonymous site conservation in the coding region of each segment was analysed using SYNPLOT2 [88] using a 25-codon sliding window and an amino-acid-based phylogenetic guide tree estimated using PhyML [89] with default parameters. Due to insertion of “NN” immediately 5′-adjacent to the splice acceptor site in segments 7 and 8 (see above), the synonymous site conservation analysis in the dual coding regions is for the M2 and NEP reading frames, respectively. To produce three-frame stop codon plots within the coding regions of each segment, the “NN”s were removed from the segment 7 and 8 sequences, and the positions of +0, +1 and +2 frame stop codons were determined in each sequence of each alignment and plotted.

Within the sequence alignments, the statistical significance of conserved open reading frames – defined as an alignment-wide absence of stop codons in a given region – was analysed for all such regions in the +1 or +2 frames that were greater than 20 codons in length. Statistical significance was evaluated by randomly shuffling zero-frame codon columns within each region and calculating the fraction of shuffled alignments that preserved an ORF in the alternative frame. 4,000 random shufflings were performed for each region. This procedure controlled for any bias for or against random long ORFs in the alternative frame that might have resulted from zero-frame amino acid use, codon use, or nucleotide biases, and also controls for phylogenetic non-independence of the aligned sequences. Four ORFs were detected with *p* < 0.05 and, for these, the number of shufflings was increased to 100,000. These regions were: segment 2, frame +2, nt 3–77, *p* = 0.0062; segment 7, frame +1, nt 689–979, *p* = 0.00017 (M2); segment 8, frame +1, nt 473–835, *p* = 0.00000 (NEP); and segment 8, frame +2, nt 12–83, *p* = 0.00085; where in all cases nt 1 corresponds to the first nucleotide of the main ORF (M1 and NS1, respectively, for segments 7 and 8), and the *p*-value is the proportion of 100,000 randomizations that have no stop codons in the given frame and region. Two of these regions are the parts of M2 and NEP that are downstream of a slice site (as indicated). The alignments contain a total of 30 regions that are > 20 codons and have no stop codons in the +1 or +2 frames. Therefore, to account for multiple testing the threshold for a 0.05 probability of a false positive was 0.05/30 = 0.00167, a threshold which is surpassed only by the M2 and NEP ORFs and by nt 12–83 of segment 8.

## Supplementary Figures

**Supplementary Figure S1:**
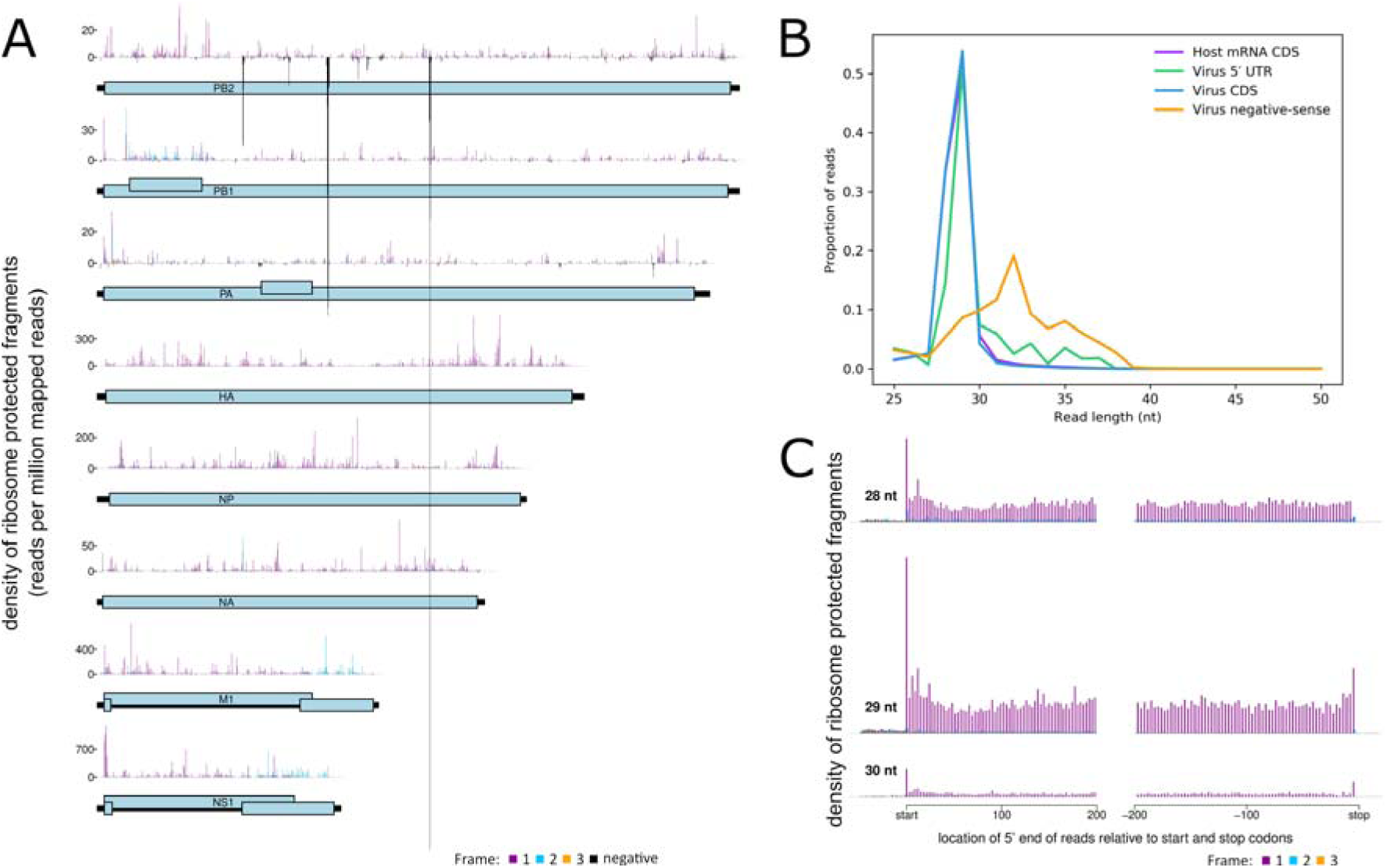
Ribosome profiling of influenza A virus mRNA. Human lung A549 cells were infected with the influenza A virus (IAV) A/Puerto Rico/8/1934(H1N1) at MOI 5. At 5 h p.i., cells were flash frozen, ribosome profiling (Ribo-Seq) was performed, and ribosome protected fragments (RPFs) were used to infer the location of ribosome P-sites on mRNAs, by mapping to the viral genome and host transcriptome. **(A)** Full Ribo-Seq profiles of all eight genomic segments of IAV (shown partially in Fig 1F), with pale blue rectangles indicating the canonical coding sequences and black rectangles indicating the genomic segments. Histograms show the location of the 5’ ends of reads with a +12 nt offset to map the approximate P-site positions. Reads which map to the first, second and third nucleotides of codons relative to the reading frame of the main ORF are indicated in purple, blue and yellow, respectively. Reads mapped to corresponding positions in the negative-sense vRNA are indicated in black on a negative scale. Within the main ORFs most reads map to the purple phase, except in the +1 frame M2 and NS2 ORFs, where most reads map to the blue phase. **(B)** Read length distributions. In preliminary work we found that IAV Ribo-Seq libraries were often contaminated by non-RPF-derived RNA which we inferred was derived from ribonucleoprotein complexes (RNPs) formed when virus nucleoprotein (NP) binds RNA. RNPs may co-sediment with ribosomes and give rise to additional nuclease-protected RNA fragments. We have found both viral and host mRNA contamination occurring at later time points of infection, suggesting that RNPs may also form with host mRNAs. High levels of contamination would make interpreting the low density of non-phased RPFs in the viral 5′ UTRs problematic. The library shown here was specifically chosen as one with relatively low contamination (assessed by a low density of reads mapping to host mRNA 3′ UTRs). To confirm that the reads observed in the viral 5′ UTRs were predominantly *bona fide* ribosome footprints, we compared their length distribution (green) with those of host mRNA coding sequence (CDS)-mapping reads (purple) and viral CDS-mapping reads (blue). The very similar length distributions indicate that the reads we saw mapping to viral 5′ UTRs in this library are mostly *bona fide* RPFs, with only a small fraction of contamination (note the high-end shoulder in the green distribution). In contrast, reads mapping to the viral genome in the negative sense orientation were found to have a very different length distribution (orange) indicating that they are, as expected, not *bona fide* RPFs, consistent with them deriving from co-sedimenting viral RNPs. **(C)** Histograms of 28, 29 and 30 nt Ribo-Seq read positions relative to annotated initiation and termination sites summed over all host mRNAs. Histograms show the location of the 5’ ends of reads with a +12 nt offset to map the approximate P-site positions. Reads which map to the 1st, 2nd or 3rd nucleotides of codons are indicated in purple, blue or yellow respectively. The vast majority of reads map to the 1st nucleotide position of codons. While only a small proportion of reads map to the 5’ UTRs, considerably fewer reads map to the 3’ UTRs.

**Supplementary Figure S2:**
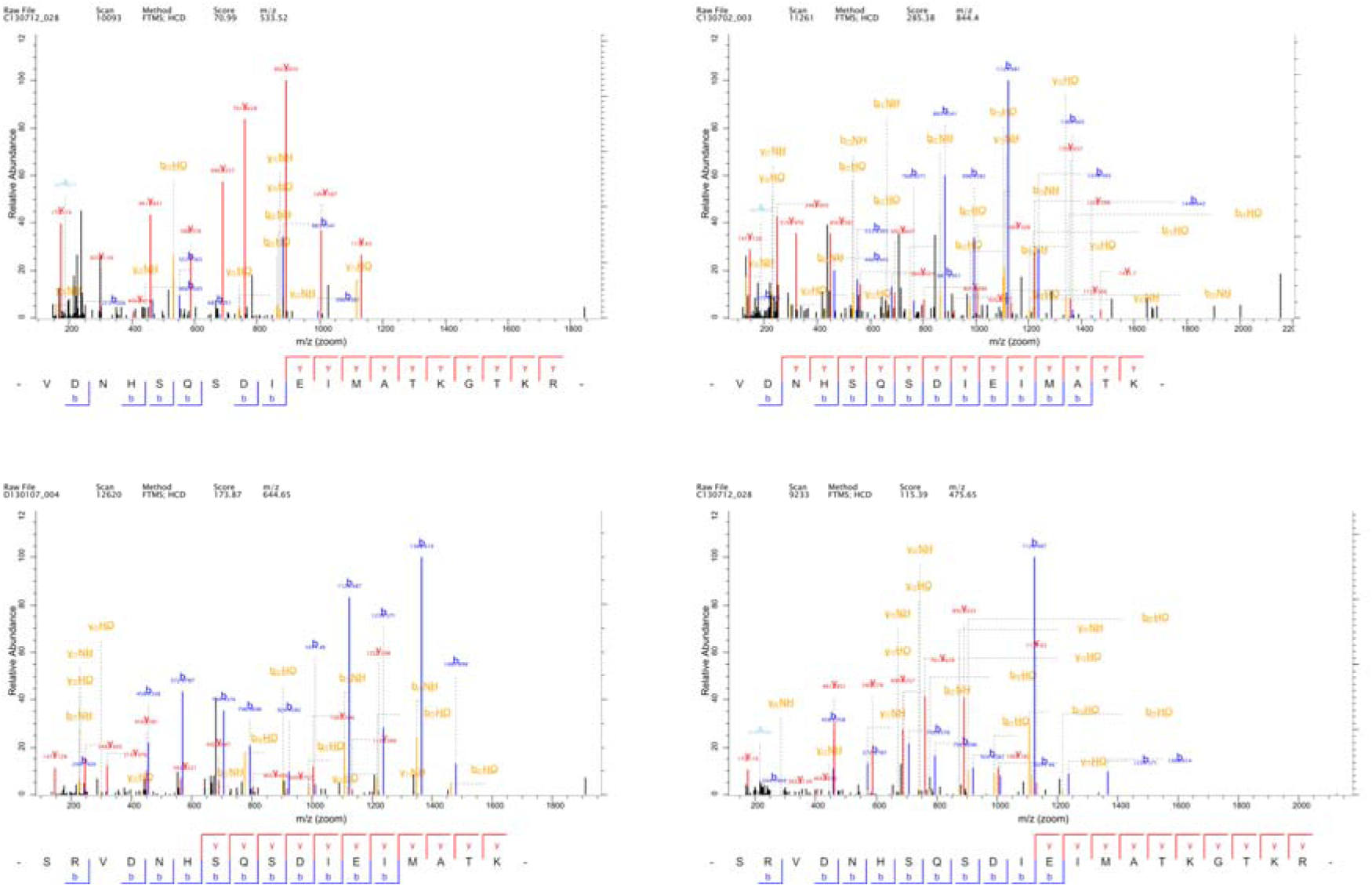
Peptide spectral matches mapping to the 5’ UTR of Influenza A Virus NP. IAV A/WSN/33(H1N1) were purified, subjected to tryptic digest and analysed by liquid chromatography and tandem mass spectrometry (LC-MS/MS). Annotated peptide spectral matches (PSMs) mapping to the 5’ UTR of IAV segment 5 (NP) are shown (see also Fig 2B and Supplementary Table S1; the canonical NP sequence begins ‘MATKGTKR…’). Four different PSMs are shown, with different numbers of missed tryptic cleavage sites (trypsin cleaves C-terminal to K or R except when followed by P).

**Supplementary Figure S3:**
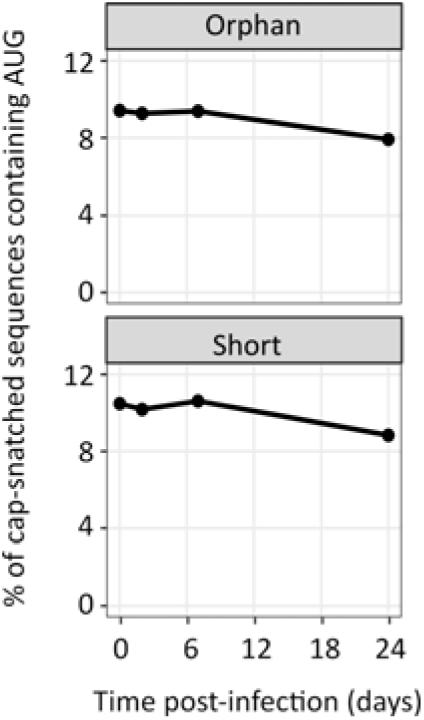
Proportion of unassigned influenza A virus cap-snatched sequences that contain an AUG. Additional data accompanying Figure 2D. Cap Analysis Gene Expression (CAGE) sequencing of cap-snatched sequences in the IAV transcriptome, collected from infected cells at various times post infection and showing the proportion of cap-snatched sequences containing AUG codons. Figure 2D describes reads that could be matched unambiguously to transcripts of a particular IAV segment; this figure describes reads where this was not possible. ‘Orphan’ reads match 3 or more nt of an IAV segment past the conserved promoter sequence, but do not align at the appropriate point, potentially due to sequencing errors or mRNA processing. ‘Short’ reads contain fewer than 3 nt of influenza sequence read past the promoter, and so cannot be unambiguously assigned to a specific genome segment.

**Supplementary Figure S4:**
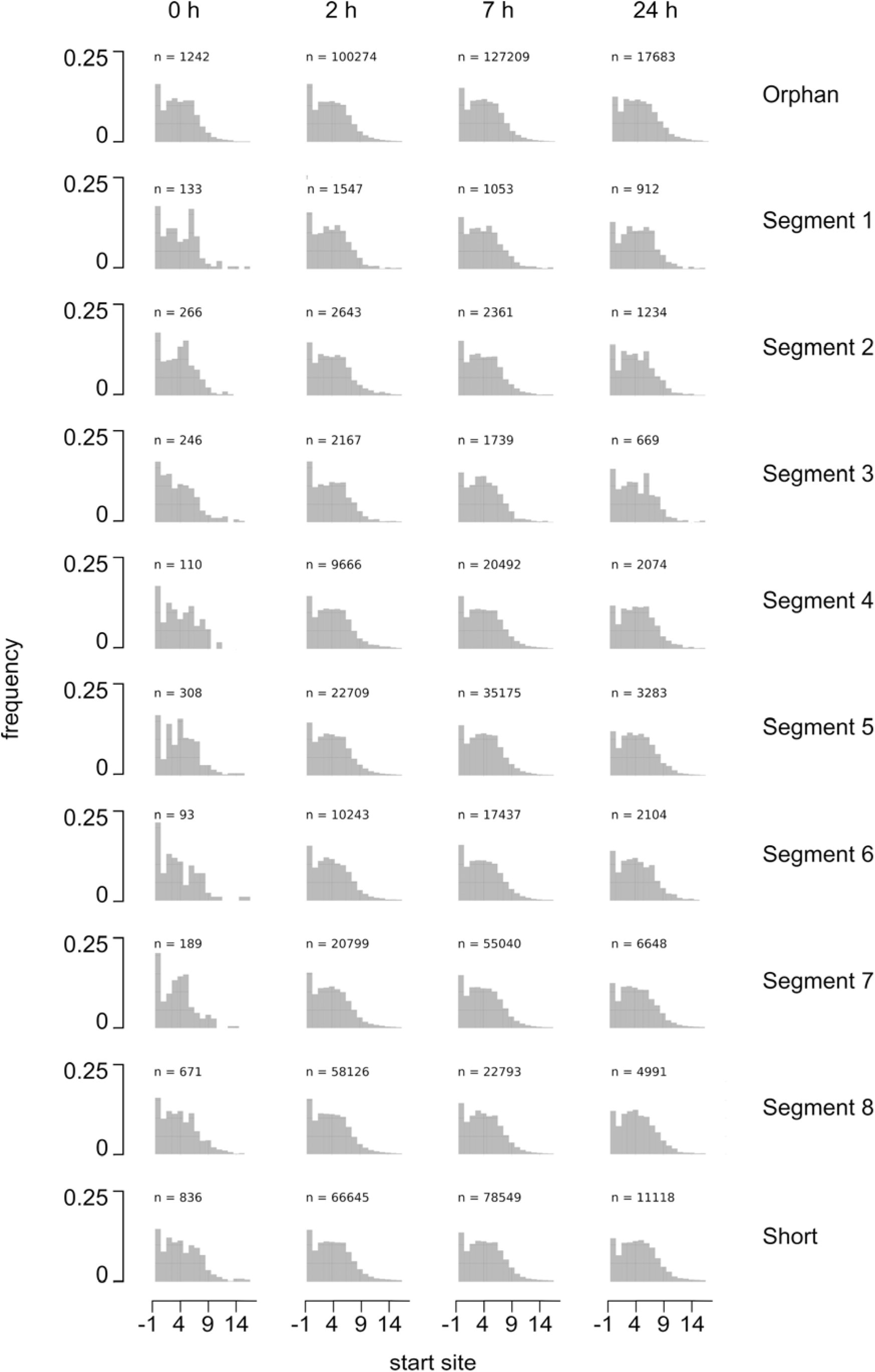
Position of AUG codons within the 5’ termini of mRNA. Monocyte-derived macrophages were infected with influenza A/Udorn/307/72(H3N2) at a multiplicity of infection (MOI) of 5 and harvested at 0, 2, 7 and 24 h post-infection. Each timepoint represents 4 donors. The sequences of viral 5’ sequences appropriated from host mRNAs by cap-snatching were determined by cap analysis of gene expression (CAGE). Where possible, reads were assigned to the viral genome segment from which the mRNA was transcribed; ‘orphan’ and ‘short’ reads are as defined in Supplementary Figure S3. Leader lengths were capped at 20 nt for analysis. The frequency of AUG codons at different positions relative to the 5’ terminus of the mRNA was expressed as a proportion of the total number of AUG-containing sequences (the number of which is indicated for each panel). Note that most leader sequences are 10 – 14 nt in length, which results in a depression in AUG frequencies at more downstream positions.

**Supplementary Figure S5:**
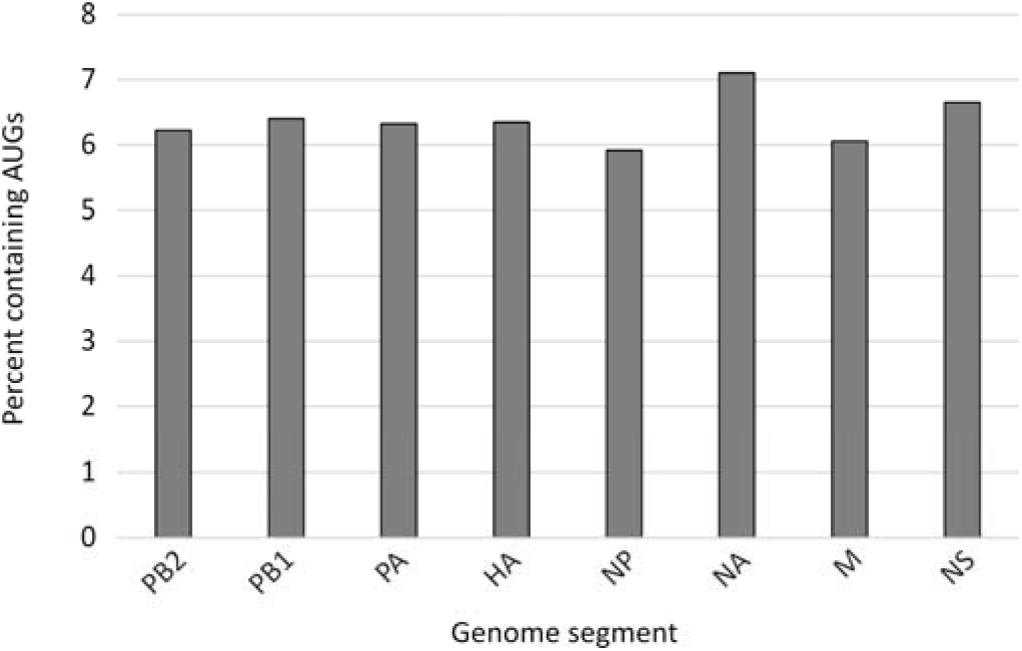
Proportion of previously-reported influenza A virus cap-snatched sequences that contain an AUG. A previously-published deep-sequencing analysis of cap-snatched leader sequences from the influenza A virus A/WSN/33(H1N1) (Koppstein *et al.* (2015) *Nucleic Acids Research* 43(10) 5052-64; doi:10.1093/nar/gkv333) was re-analysed. The percentages of cap-snatched leader sequences from each genome segment that contain AUG codons are shown.

**Supplementary Figure S6:**
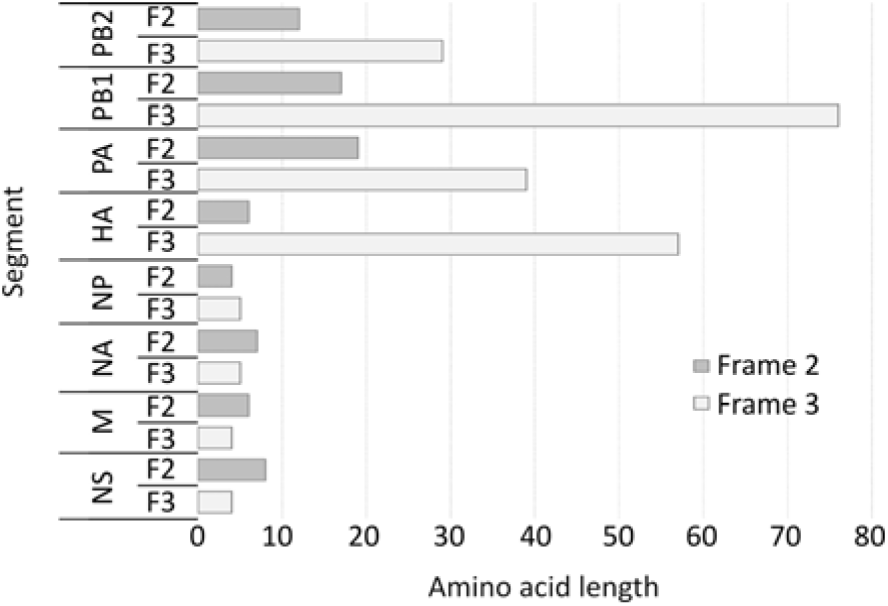
Overlapping uORFs in the influenza A virus maCa04. The lengths of 5’ ORFs that overlap the canonical ORFs (frame 1) of the mouse-adapted influenza A virus ma-A/California/04/2009(H1N1). The lengths shown are the number of codons that can be read from the 5’ end of each segment (coding sense, including the 5′ UTR) in frames 2 and 3 before encountering a stop codon.

**Supplementary Figure S7:**
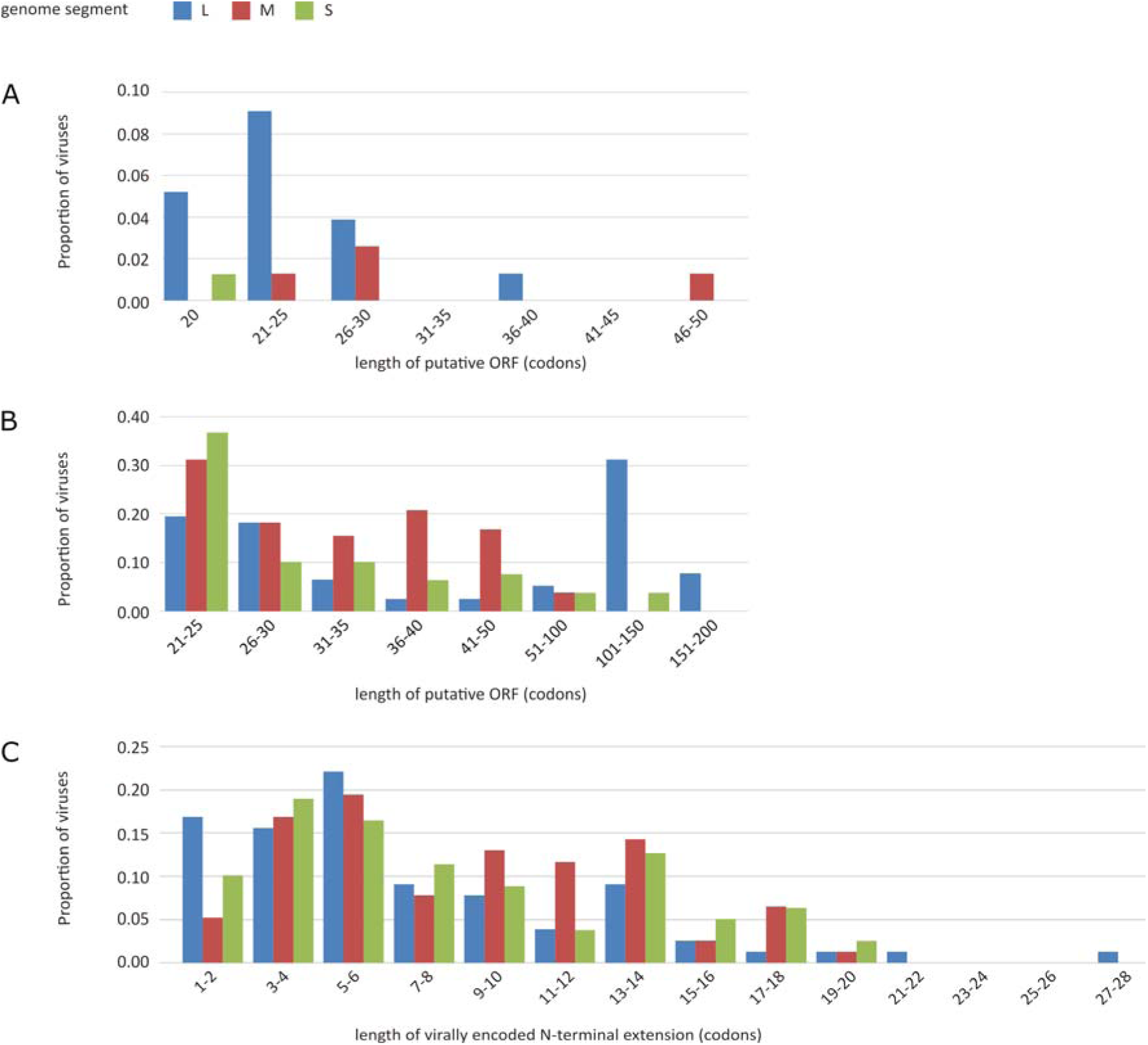
Putative upstream ORFs in peribunyaviruses. Full-length L, M and S genome segments from 78 different peribunyavirus species were translated in all three forward frames, numbered with respect to the main ORF of that segment. The lengths of putative uORFs are shown, defined as sequences that could be read from within the 5’ UTR of a segment **(A)** in any frame, continuing for more than 20 codons before encountering a stop codon upstream of the canonical ORF; **(B)** in frames +1 and +2 with respect to the canonical ORF, overlapping the canonical ORF and continuing for more than 20 codons before encountering a stop codon; and **(C)** in frame with the canonical ORF and encoding a potential N-terminal fusion to that ORF, of any length.

**Supplementary Figure S8:**
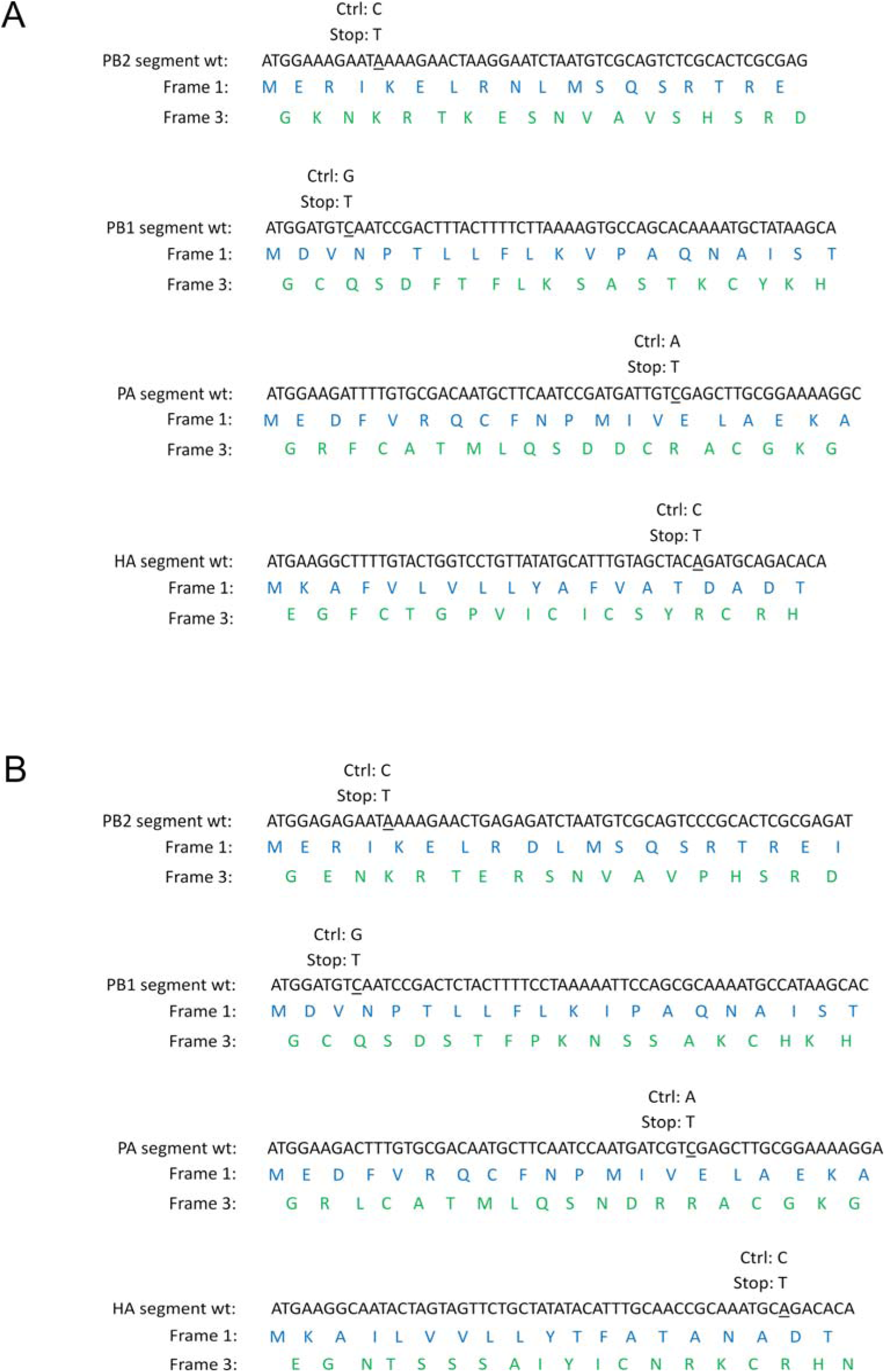
Design of frame 3 mutations in influenza A viruses. Partial nucleotide and amino acid (frames 1 (canonical ORF) and 3) sequences for the PB2, PB1, PA and HA segments of influenza A viruses, beginning at the canonical start codon. Annotations at mutated sites indicate WT sequences (underlined) and mutations designed to introduce frame 3 stop codons (‘Stop’) or to be conservative in frame 3 (‘Ctrl’). All mutations are synonymous in frame 1. Designs for mutants are shown for **(A)** influenza A/WSN/33(H1N1) and **(B)** mouse-adapted influenza A/California/04/2009(H1N1).

**Supplementary Figure S9:**
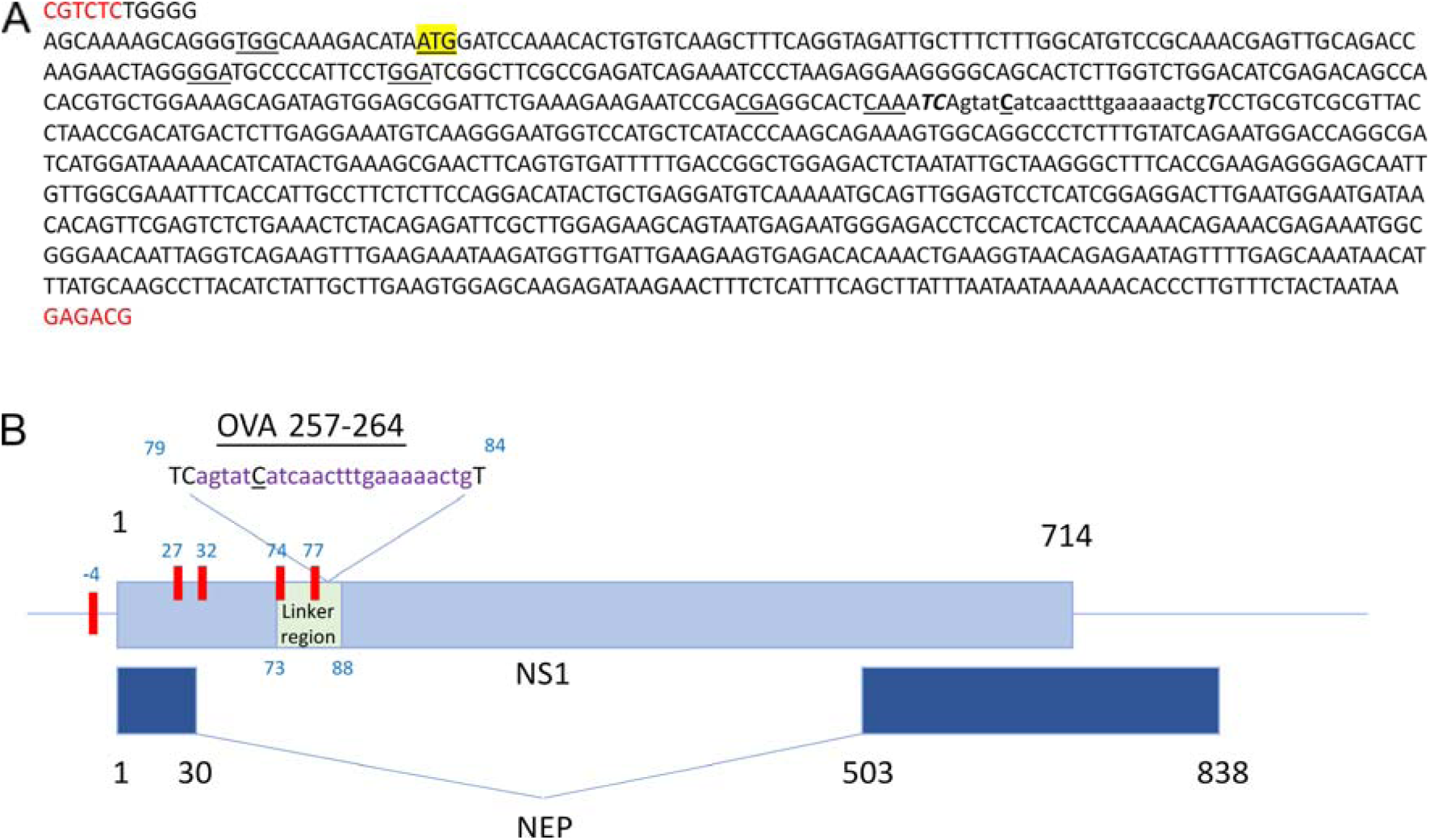
Design of PR8-NS.F3.SIIN virus. Mutagenesis design for inserting the OVA_257-264_ (SIINFEKL) epitope into frame 3 of segment 8 of the influenza A virus genome, in a region corresponding to the linker sequence of the NS1 protein (encoded in frame 1). **(A)** A synthetic sequence designed for cloning into the plasmid pDUAL:NS, which encodes segment 8 of the influenza A/Puerto Rico/8/1934(H1N1) (PR8) virus backbone. BsmB1 restriction sites are indicated in red, and the start codon of NS1 and NEP is highlighted in yellow. A sequence encoding SIINFEKL replaced codons 79 – 84 of NS1 (lowercase italic). The replacement sequence was flanked by two upstream nucleotides and one downstream nucleotide to introduce a frameshift into frame 3; it was also subject to an additional synonymous mutation to remove a stop codon in frame 1 (uppercase bold italic). Frame 3 stop codons upstream of the replacement sequence (underlined) were eliminated by point mutations that were synonymous in frame 1. **(B)** In a reverse genetic experiment, PR8 virus was generated using this sequence. The organisation of segment 8 of this virus is shown schematically, with blue boxes indicating ORFs, a horizontal line indicating the UTRs of frame 1, and red boxes indicating mutated frame 3 stop codons. Numbering of codon positions is with respect to the NS1 and NEP start codon.

**Supplementary Figure S10:**
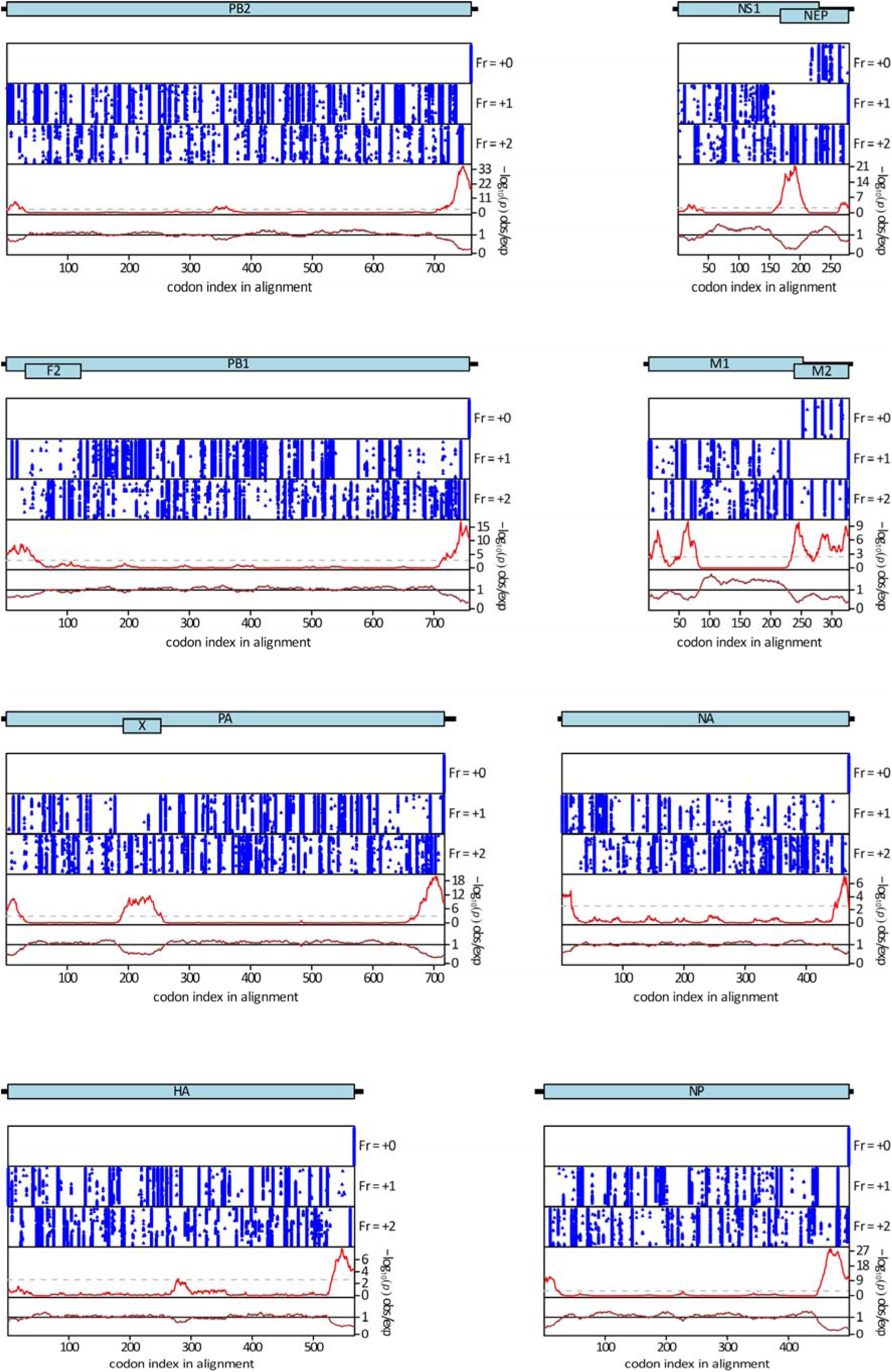
The eight segments of the influenza A virus genome are shown as schematics, with pale blue rectangles indicating the coding sequences and black rectangles indicating the UTRs. Below each segment map, the upper three panels show the positions of stop codons (blue dots) in each of the three forward reading frames in each of the aligned sequences, within the coding regions of the segment. The lower two panels show the synonymous site conservation analysis: the red line shows the probability that the observed conservation could occur under a null model of neutral evolution at synonymous sites (with a dashed line indicating p = 0.05 after multiple testing correction) and the brown line depicts the ratio of the observed number of substitutions to the number expected under the null model.

## Supplementary Tables

Supplementary Table S1: Peptide spectral matches mapping to the 5’UTR of influenza A virus NP

Supplementary Table S2: Peribunyavirus sequences consulted

Supplementary Table S3: Details of putative uORFs in peribunyavirus L segments

Supplementary Table S4: Details of putative uORFs in peribunyavirus M segments

Supplementary Table S5: Details of putative uORFs in peribunyavirus S segments

Supplementary Tables S1 – S5 are provided in the Excel spreadsheet Sloan_sNSVuORFS_Supplementary_Tables.xlsx

## Acknowledgements

The authors would like to thank Ervin Fodor, University of Oxford for support and critical comments on the project; Svenja Hester, Benjamin Thomas and Shabaz Mohammed of the Advanced Proteomics Facility, University of Oxford for proteomics; Elly Gaunt, University of Edinburgh for critical reading of the manuscript; and staff within the Institute of Infection, Immunity and Inflammation Flow Cytometry Facility, the Central Research Facility at the University of Glasgow, Thomas Purnell of the Institute of Infection, Immunity and Inflammation, University of Glasgow and Dimitris Athineos of the Beatson Institute for technical assistance and discussion.

